# K-Ras G-domain binding with signaling lipid phosphoinositides: PIP2 association, orientation, function

**DOI:** 10.1101/324210

**Authors:** Shufen Cao, Stacey Chung, SoonJeung Kim, Zhenlu Li, Danny Manor, Matthias Buck

## Abstract

Ras genes are potent drivers of human cancers, with mutated K-Ras4B being the most abundant isoform. Targeted inhibition of oncogenic gene products is considered the holy grail of present-day cancer therapy, and recent discoveries of small molecule inhibitors for K-Ras4B greatly benefited from a deeper understanding of the protein structure and dynamics of the GTPase. Since interactions with biological membranes are key for Ras function, the details of Ras - lipid interactions have become a major focus of study, especially since it is becoming clear that such interactions not only involve the Ras C-terminus for lipid anchoring, but also the G-protein domain. Here we investigated the interaction between K-Ras4B with the signaling lipid phosphatidyl inositol (4,5) phosphate (PIP2) using NMR spectroscopy and molecular dynamics simulations, complemented by biophysical and cell biology assays. We discovered that the β2 and β3 strands as well as helices 4 and 5 of the GTPase G-domain bind to PIP2, and that these secondary structural elements employ specific residues for these interactions. These likely occur in two orientation states of the protein relative to the membrane. Importantly, we found that some of these residues, which are known to be oncogenic when mutated (D47K, D92N, K104M and D126N), are critical for K-Ras-mediated transformation of fibroblast cells, while not substantially affecting basal and assisted nucleotide hydrolysis and exchange. We further showed that mutation K104M can indeed abolish localization of mutant K-Ras to the plasma membrane. These findings suggest that specific G-domain residues play an important, previously-unknown role in regulating Ras function by mediating interactions with membrane PIP2 lipids. Thus, a detailed description of the novel K-Ras-PIP2 binding surfaces is likely to inform the future design of therapeutic reagents.

## Introduction

Ras GTPases regulate diverse signal transduction pathways, controlling cell proliferation, differentiation and apoptosis, organization of the cytoskeleton, vesicular transport, metabolism, and nuclear import [e.g. Karnoub et al., 2008; Pylayeva-Gupta et al., 2011]. Mutated Ras proteins are associated with approximately 30% of all human cancers, where they drive oncogenic processes [Malumbres et al., 2003]. And among the three main isoforms expressed human cells - N-Ras, H-Ras, and K-Ras [Cox et al., 2010], mutated K-Ras4B is the most abundant of the oncogenic GTPases [Prior et al., 2012; Cox et al., 2014]. The protein is comprised of two major domains: the G-domain and the hyper-variable region (HVR) (Fig. 1A). The G-domain includes the N-terminal residues 1-165, which associate with GTPase exchange factors (GEFs), and GTPase activating proteins (GAPs) [Vetter et al., 2001].

**Figure 1.**
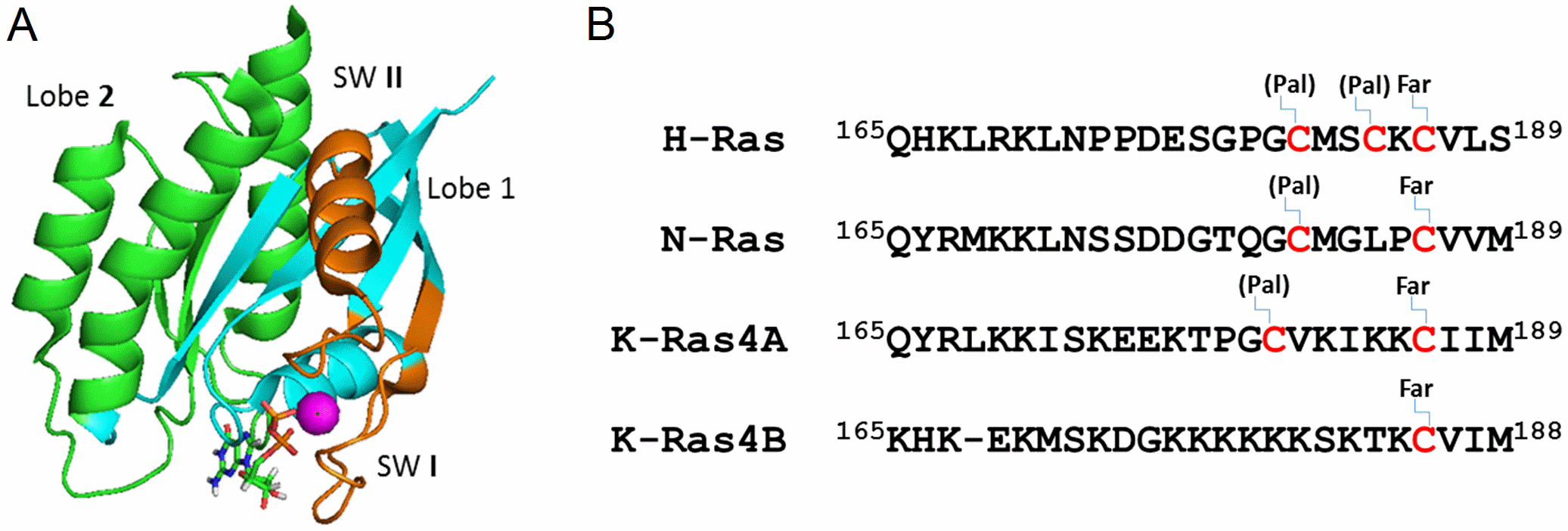
Structure of K-Ras. A. Structure of K-Ras4B (G12V, 1-169) is shown in ribbon representation with lobes, nucleotide and switch regions highlighted as follows: Lobe 1 and 2, as well as the switch regions are shown in cyan, green and dark orange, respectively. The bound GTP is shown in stick representation and the Mg^2+^ ion as a cyan sphere. B. Sequence alignment of the HVR domains of K-Ras, H-Ras and N-Ras, with posttranslational lipid modifications, but not carboxy-methyl truncations indicated. Pal and Far refer to palmitoylation and farnesylation sites, respectively.

The G-domain is comprised of two lobes (Fig. 1A). Lobe 1, the catalytic subdomain, encompassing residues 1-86, contains the functionally critical switch regions whose conformation and dynamics is nucleotide dependent (switch1 including residues 25-40, and switch 2 including residues 57-75), as well as the phosphate binding region (P-loop, residues 10-17). The second lobe, comprised of residues 87-166, is a regulatory domain and contains allosteric regions such as helix 3 and loop 7. Intramolecular communication has been proposed between the allosteric domain and the catalytic domain at the other side of the protein [reviewed in Gorfe et al., 2010; Nussinov et al., 2013]. As found in all Ras GTPases, the HVR is located at the end of the second lobe, in K-Ras4B comprising a 24 residue segment with the very C-terminal CAAX sequence (C, cysteine; A: aliphatic amino acid, X: any amino acid), which undergoes farnesylation, truncation and methylation and serves as a lipid anchor to the plasma membrane. In contrast to the high sequence identity in the G-domain, Ras isoforms differ significantly in the HVR and can undergo isoform specific post-translational lipid modification (Fig. 1B). The K-Ras4B HVR undergoes a single farnesylation, whereas some other GTPases can utilize an additional palmitoylation for further anchoring to membranes. The unique C-terminal polybasic region further aids in localizing K-Ras4B to lipid bilayers [Wright et al., 2006; Brunsveld et al., 2009; Chenette et al., 2011].

It has long been established that interactions with cellular membranes are critical for signaling by Ras GTPases. The major outcome of these interactions is thought to be a temporal and spatial sequestration of a multi-component signaling complex in the proximity of an activated growth factor receptor [Ahearn et al., 2011]. While the majority of the research in Ras GTPase interaction with membrane has focused on its phosphatidylcholine and phosphatidylserine components, the interaction of Ras GTPases with phosphoinositides have been rarely studied.

Of the signaling lipid classes, phosphoinositides regulate key aspects of cell growth and proliferation. For example, PIP(4,5)P2 plays a critical role in endosomal vesicle trafficking from an to the apical as well as basolateral plasma membrane, assembly of the actin cytoskeleton, and communication with the extracellular milieu, whereas PI(3,5)P2 is responsible for late endosomal trafficking [Di Paolo et al., 2006; Yang et al., 2018]. Under pathological conditions, phosphoinositides serve as key mediators of aberrant proliferation and survival signals, rendering them important targets for therapeutic interventions [Wymann et al., 2008]. In the context of cancer, phosphoinositides were shown to be major regulators of cell motility, invasion, and metastasis [Yamaguchi et al., 2010; Gagliardi et al., 2015; Lien et al., 2017]. Recent studies revealed, several classes of transmembrane receptors and ion channels might be regulated by PIP2, or at least bind this signaling lipid with considerable affinity [Hedger et al., 2015; Kolay et al., 2016, Chadli et al., 2017]. Thus, PIP2 signaling may results in a cell-global concerted response to environmental changes.

It is, therefore, likely that localization if not activity of Ras GTPases is also affected by changes in nearby levels of PIP2. Furthermore, PI3K is a major effector of Ras, which when activated by the GTPase generates PIP3 from PIP2(4,5) [Yang et al., 2012], while PLCβ and γ are regulated by Ras to break down PIP2 [Gresset et al., 2012]. Potentially, PIP2 and Ras could be involved in positive or negative regulatory feedback loops. PIP2 is known to play a role in K-Ras localization in cells [Heo et al., 2006; Gulyas et al., 2017] but the mechanism is debated and remains unclear [e.g. Zhou and Hancock, 2017]. At the protein level, we have the opportunity to understand the molecular (structural/dynamics) details of the interactions, and especially in the context of HVR isoform- and compartment-specific Ras signaling those rules are just beginning to be uncovered. For example, a number of studies have well established that the dynamic interactions of the Ras HVR motifs with membranes modulate the targeting of Ras to cell surface and intracellular organelles [Magee et al., 1987, Baker et al., 2003, Rocks et al., 2005]. In addition, different HVR motifs in Ras isoforms also specify localization within plasma membrane subdomains [Prior et al., 2001, Prior et al., 2005] and recently the Hancock laboratory showed how mutations in the K-Ras4B HVR can switch phosphotidylserine to PIP2 mediated membrane localization [Zhou et al., 2017].

However, we and others have substantiated the view that the Ras G-domain - lipid interactions can also play a significant role in determining the protein’s configuration (i.e. G-domain orientation) and dynamics at the membrane. For example, recent computational investigations suggested the participation of K-Ras G12V G-domain in binding with POPS membrane [Gorfe et al., 2007b; Prakash et al., 2016, Li et al., 2016; Li et al., 2017] and with PIP2 containing membranes [Li et al, 2017; Gregory et al., 2017]. Experimental work by Ikura and colleagues [Mazhab-Jafari et al., 2015] also provided evidence that the K-Ras G-domain exists in several distinct orientations when contacting a bilayer of POPS lipid membrane. However, a detailed experimental study on K-Ras4B ‒ PIP2 interactions has been missing.

Here, we utilized in vitro binding assays, NMR spectroscopy, computational simulations and cell function experiments to study the K-Ras interaction with a PIP2 containing membrane. Our data clearly demonstrate a direct interaction between K-Ras4B and PIP2, also of the GTPases’s G-domain and we provide structural details of the interactions and of the K-Ras4B orientations relative to the lipid bilayer. Lastly, the functional relevance of K-Ras4B-PIP2 interactions are corroborated when key residues are mutated and such PIP2 binding compromised GTPases are examined in cell-transformation activity and intracellular localization experiments. Implications of these findings for Ras localization and function in cell signaling are discussed.

## Results

### Full length as well as HVR-truncated K-Ras4B binds to PIP2 and other specific lipids

Lipid strip assays were first carried out to screen the lipids that can bind to unmodified K-Ras4B (the HVR is not lipidated or tuncated, as we are interested in the initial membrane interactions that the Ras G-domain may make). Fig. 2 shows the results for the binding of full length (top) and HVR-truncated (bottom) K-Ras4B with different lipids. It can be seen that full length K-Ras4B can bind to LPA, singly phosphorylated inositol phosphates, PIP2, PA and PS. Comparing the results of full length and truncated K-Ras, overall, lipid binding with the full length K-Ras is stronger. Additionally, when the HVR domain was truncated from K-Ras4B, the interactions with some lipids were abolished (e.g. PIP3, LPA, PS with truncated G12V.GMPPNP K-Ras as well as truncated S17N.GDP K-Ras4B). These observations suggest that the HVR of K-Ras is necessary for binding of these lipids. However, binding with singly phosphorylated inositol phosphates, with PI(3,4)P2, PI(3,5)P2, PI(4,5)P2 are maintained for HVR truncated K-Ras proteins both in GMPPNP and S17N GDP forms, although the binding with PIP2s is clearly weaker, compared with the full length protein. Following our presentation of these data at the NCI Ras conference in early Dec. 2015, Nussinov and Gaponenko showed a lipid strip assay of GDP loaded K-Ras4B in a Current Opinions Review (Banerjee et al., 2016). This overall is consistent with our data showing that PA binding is associated mainly with the full length protein and possibly with the G12V mutation in the truncated protein. In summary, our observations show that the K-Ras4B G-domain binds with selective lipids.

**Figure 2.**
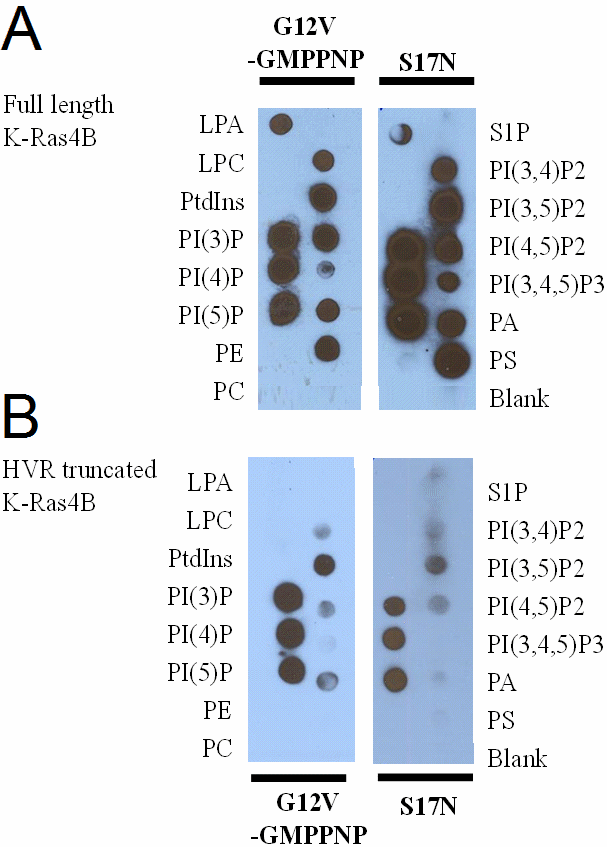
Lipid strip assay shows binding and that the interactions are lipid, HVR but less so GTPase-state selective. A. full length K-Ras4B (1-188); and B. HVR truncated K-Ras4B (1-169) as i) G12V constitutively active mutant loaded with GMPPNP, ii) dominant negative mutant S17N, bound to GDP.

### K-Ras4B binds to PI(4,5)P2 moderately well

Our lipid strip assays showed that K-Ras4B can interact with PIP2s including PI(3,4)P2, PI(3,5)P2, PI(4,5)P2. This study specifically focuses on the binding between K-Ras4B with signaling lipid PI(4,5)P2. To further confirm the binding between K-Ras4B and PI(4,5)P2, microscale thermophoresis (MST) was employed, also for measuring the binding affinity. The purified K-Ras4B protein was fluorescently labeled with dye NT-647, and then was titrated against PIP2 doped liposomes (5% PIP2, 95% DOPC). A representative datasets for full length unmodified and HVR truncated K-Ras4B are shown in Fig. 3, and the fitted Kd is 28 ± 7 μM, indicating a moderate binding between K-Ras4B and PI(4,5)P2 in vitro. However, the binding of HVR truncated K-Ras to these liposomes is significantly decreased (about 9 fold) pointing to the importance of the HVR to enhance affinity.

**Figure 3.**
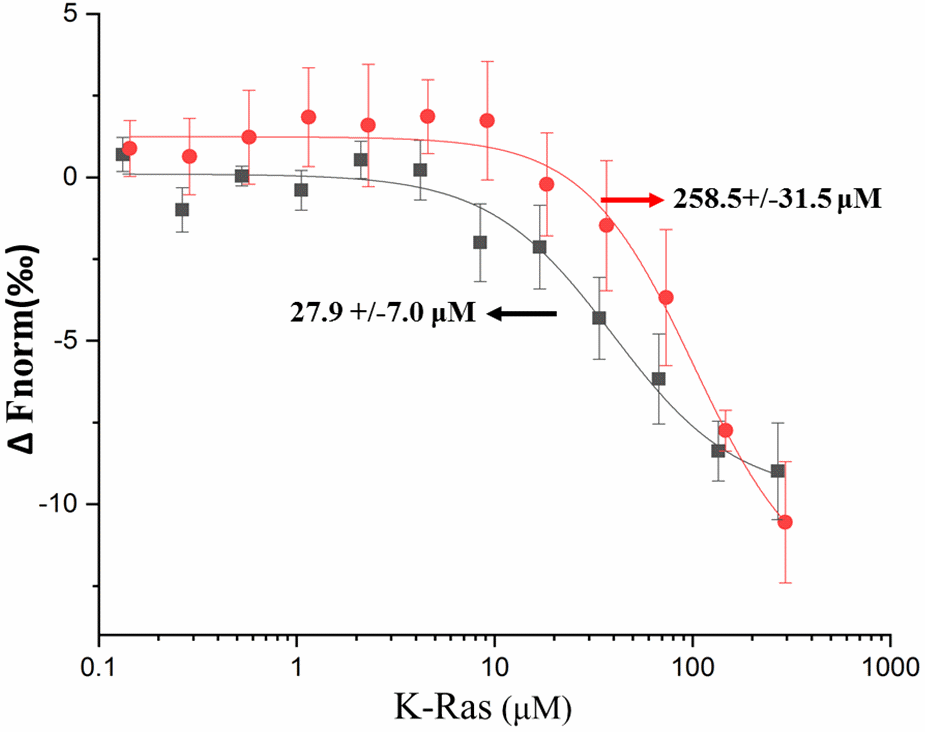
MST measurement of full length K-Ras4B and HVR truncated K-Ras4B interaction with 5% PI(4,5)P2 doped DOPC lipid vesicles. A DOPC liposome doped with fluorescently labeled and unlabeled PI(4,5)P2 was titrated with a serial dilution of full length K-Ras4B (1-188) G12V.GMPPNP or HVR truncated K-Ras4B (1-169) G12V.GMPPNP in a twofold dilution series.

### K-Ras4B on PI(4,5)P2 membrane – interface and orientations studied by NMR

The NMR spectrum of HVR truncated K-Ras4B has been assigned (Vo et al., 2013.) and we first sought to characterize PIP2 lipid binding to the K-Ras4B G-domain by use of the lipid headgroup, IP3. However, the perturbations to the NMR spectra were both small and distributed across the protein, making it unlikely that all changes reflect the true configurational states at a lipid bilayer surface (see SI Fig.1 and Table S1). Recent developments in NMR have made it possible to study integral membrane proteins using lipid bilayer-like discs, bicelles or nanodiscs, up to 10 nm in diameter [Prosser et al., 2006, Glück et al., 2009]. Here, we next used PIP2 doped nanodiscs. Slightly smaller perturbations were seen compared to the lipid head-group, overall affecting a similar number of resonances and the data was not analyzed further. It is possible that nanodiscs are too small to allow good diffusion of the lipid spin label (approx. 2 per disc), which may on average also occupy a non-interacting side. However, increasing the spin label concentration did not improve the discrimination between interacting and non-interacting residues, suggesting, as an alternative that K-Ras may interact non-specifically and transiently with the disc-bounding peptide in absence of membrane anchoring. Thus, we decided to go to an even bigger membrane model system, LUVs (large unilamellar vesicles) or liposomes.

NMR spectroscopy is a unique method for the characterization of weak and dynamic interactions [Vinogradova et al., 2012; Barrett et al., 2013]. As mentioned by contrast to the studies of Ikura and colleagues, do not have an anchoring of K-Ras4B via a lipidated C-terminus, which would lead to more persistent membrane contacts and, even with a nanodisc, a corresponding increase in the molecular correlation time, resulting in an extensive linebroadening, especially of the backbone resonances [Mazhab-Jafari et al., 2015]. In our case interactions are very transient to the extent that with a liposome (of 100 nm in diameter) effects due to proximity of a spin label such as paramagnetic ion gadolinium (Gd3+), far outweigh the effects due to transient attachment and large correlation times. In such a context, PRE appears to be a sensitive tool for transient interactions [Ubbink et al., 2001, Mazhab-Jafari MT et al., 2013 and also Clore et al., 2015 as well as Ceccon et al., 2016].

Fig. 4A shows the superimposed ^15^N-^1^H HSQC-TROSY spectra of full length K-Ras4B.G12V.GMPPNP with PIP2 doped DOPC lipid vesicles without (red) and with (green) Gd^3+^ spin-labeled lipid. A decrease in peak intensity indicates proximity to the spin labeled lipid, taken as binding to the liposome. Fig. 4B is a plot of the peak intensity ratio versus protein sequence (only to res. 159, as the remainder has not been assigned. Many signals in the center of the spectrum, fully shown in Fig. S3, overlap and, thus, are not clearly attributable, with some likely belonging to the HVR). The intensity of a good number of NMR signals from the amide groups in the G-domain are perturbed, indicating the involvement of these residues in binding which brings them close to the spin label. Specifically, the perturbations are mainly in the catalytic lobe and fewer in the allosteric lobe. In the catalytic lobe, the perturbations are mostly located at the strands β1, β2 and β3, the C terminus of α3. In the allosteric lobe, the perturbations are mainly at the termini of β6 and the loop between β6 and helix α5, occasionally at helix α3. **Supplementary table 1** lists the residues that are implicated by these data to interact with the PIP2(4,5) doped liposome. Those residues were mapped to the 3D structure of K-Ras4B G-domain, as shown in Fig. 4C. The distribution of these residues suggests that not all residues involved in interaction can be simultaneously satisfied by a single protein-lipid interface. This indicates a dynamic equilibrium between at least two orientations of K-Ras4B on the PIP2 membrane, employing different sites of K-Ras for membrane association.

**Figure 4:**
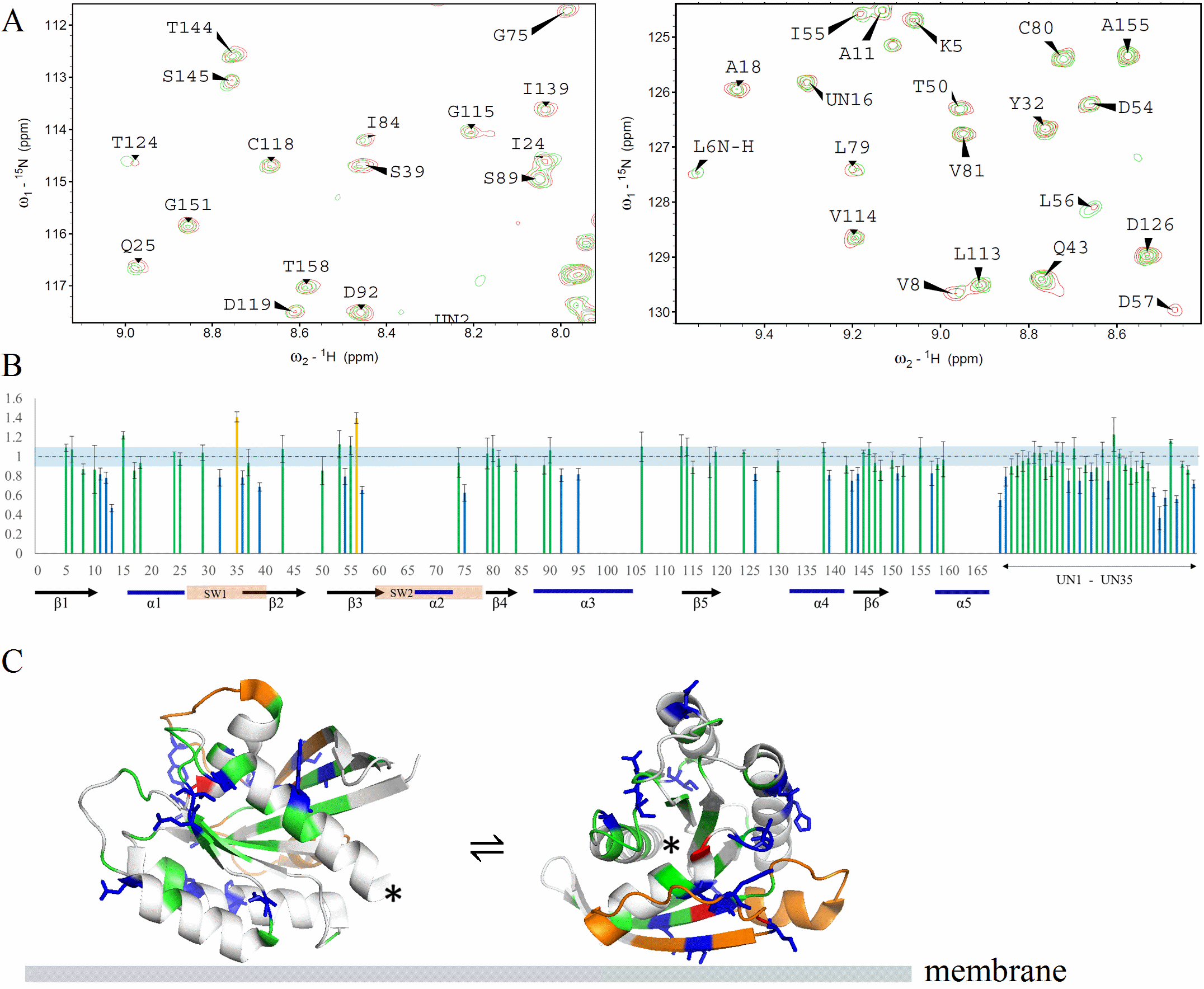
Full length K-Ras4B (1-188) interaction with PIP2 doped DOPC liposomes studied by NMR spectroscopy. A. Representative areas of superimposed ^15^N-^1^H HSQC-TROSY spectra of full length K-Ras4B.G12V.GMPPNP recorded at 800 MHz and 37 °C, in the presence of PIP2 doped DOPC lipid vesicles without (red) and with (green) Gd3+ spin-labeled lipid (The full range spectrum is shown in supplementary Fig. 3). B. Peak intensity change as a function of protein sequence. The residues decreased >10% in peak intensity are shown as blue bars, the residues increased >15% in peak intensity are shown as red bars, and the residues changed within ± 15% are shown as green bars. Residues positions not assigned in the NMR spectrum are left blank, but the data are shown as UN1 to UN27 at the end of the sequence. The K-Ras4B switch regions as well as the secondary structure are indicated at the bottom. C. Peak intensity changes are mapped to the 3D structure of K-Ras4B.G12V (PDB: 4TQ9) and are displayed as two orientations of the protein. The color scheme is same as b), with unassigned residues shown in grey, and the switch regions shown in orange (the location of the C-terminus is indicated by a *).

We further investigated the interaction of HVR truncated K-Ras with the PIP2 doped liposome. Fig. 5A shows the plots of the peak intensity ratio versus protein sequence. Several residues in the G-domain still experience diminished peak intensity, confirming the interaction between truncated K-Ras4B and PIP2 liposome. The perturbed residues in truncated K-Ras include part of the β2-and β3-strands in the catalytic lobe. Similar to the full length protein, peaks from residues located in β6 and in the turn between β6 and helix α5 experience the most significant intensity changes, as well as for two unassigned residues. A mapping of perturbed residues onto the 3D K-Ras structure is shown in Fig. 5B. However, the extent of intensity change is diminished compared to the full length K-Ras, consistent with the other experiments above, which showed that binding is more transient for HVR truncated K-Ras. Furthermore, the perturbed residues are much less spread out in the protein sequence, possibly because the HVR-lipid interaction also creates longer range effects in the full length protein. Nevertheless, again the distribution of these residues in the structure is better accommodated by two or more orientations of K-Ras4B on membrane than by a single orientation.

**Figure 5:**
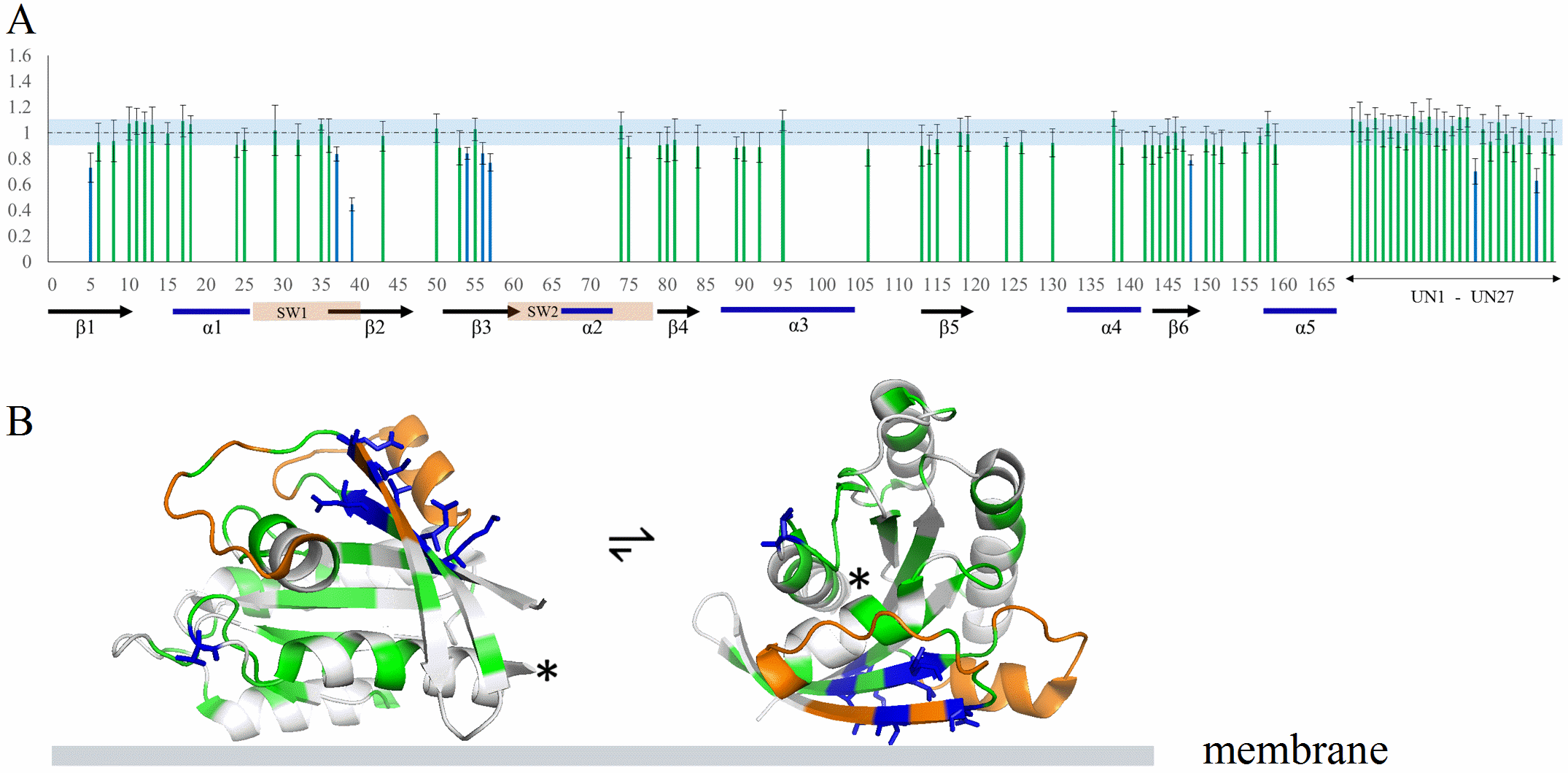
Truncated K-Ras.G12V.GMPPNP (1-169) interaction with PIP2 doped DOPC liposomes studied by NMR spectroscopy. The legend to A and B, is the same as for Fig. 4 B and C, respectively.

### K-Ras4B on PI(4,5)P2 membrane - interface and orientations studied by simulation

All-atom classical MD simulations were also used to examine the K-Ras4B regions that interact with PIP2. Because the re-orientation of the K-Ras4A GTPase at membranes is slow [Li et al., 2017], simulations were started with two orientations of K-Ras4B (orientation states OS1 and OS2) previously predicted by more extensive simulations at a POPS containing bilayer [Prakash et al., 2016b]. Then simulations of K-Ras4B interacting with a PIP2 containing membrane were performed for 380ns, sufficient to allow clustering of PIP2 near the protein. The results show that the two orientations have remained stable over the course of each of the trajectories. It should be noted that for our previous K-Ras4A [Li et al., 2017] and K-Ras4B [Gregory et al., 2017] simulations with PIP2, primarily the OS1 state was found to be populated (similar to O3 in Li et al., 2017). The OS2 started simulation may not be able to interconvert to OS1 on the timescale of the current simulations, also having been stabilized by PIP2 contacts; such transition was seen, however, by Gregory et al. using a membrane model with increased fluidity.

Snapshots for the final configurations of K-Ras4B with respect to the membrane are shown in Fig. 6A. In both orientations, the G-domain as well as the HVR domain contact the membrane. But the two orientations involve different regions of the G-domain. In the OS1 orientation, the interaction mainly involves α-helices 3 and 4 in the globular domain, while the OS2 orientation primarily involves the β-sheet and α-helix 2. Fig. 6B shows the specific residues which frequently contact the membrane during the simulation. The HVR region in both cases involves large number of contacts with the membrane, due to the insertion of farnesyl group into the membrane as well as the favorable interaction between the multiple lysine groups (stretch ^175^KKKKKKSKTK^184^ of the HVR) and the membrane. In the G-domain, the membrane contacting residues are mostly positively charged residues. But several negatively charged residues also participate in interactions with the membrane, such as E3, E107 and D132. While the positively charged residues directly interact with PIP2 phosphate groups in membrane, the interaction between negatively charged residues and the PIP2 phosphate was revealed to be bridged by the Na+ ions, as shown in Fig. 6C, with representative interactions between Arg/Lys/Glu residues and PIP2 lipid. Overall, the electrostatic interactions likely play a major role in K-Ras interaction with the PIP2 in the membrane. Moreover, residues showing high contact frequency with the membrane in simulations are consistent with the NMR experimental results, not only the exact same residues but also close by residues in the experiments (also considering gaps in the experimental data since less than half of residues are confidently assigned). Examples are K5 in β1 (K5 in expts.), R41 in β2 (S39 in expts.), R97 in α3 (H95 in expts.), K128 in α4 (D126 in expts.), S136, Y137, G138 in α4 (I139 in expts.)

**Figure 6.**
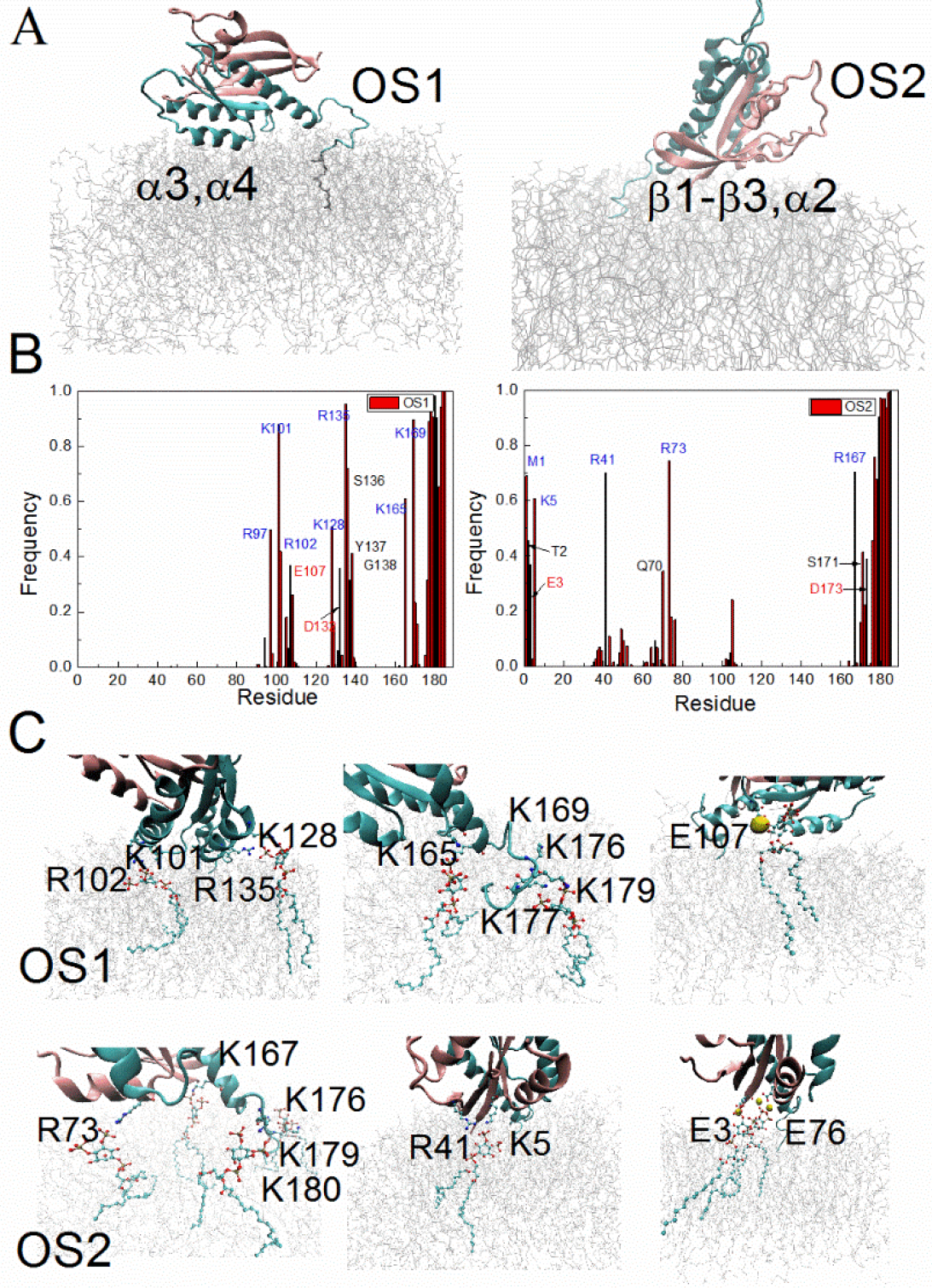
Simulation of K-Ras4B at a membrane composed of POPC and PIP2. A. Snapshots of the orientation of K-Ras4B at the PIP2 containing membrane at the end of simulation; left, OS1- and right, OS2-started simulation. B. Contact frequency of K-Ras4B residues interacting with the PIP2 membrane (residues within 3 Å of the membrane surface are considered). Top: OS1 orientation, Bottom: OS2 orientation. C. Representatives interactions between Arg/Lys/Glu residues and PIP2 lipids in each of the final structures.

### The effect of mutations on biological function: excluding an effect on intrinsic and regulated GAP and GEF activity of K-Ras4B by use of in vitro assays

The NMR and simulations data above suggest a number of surface clusters of residues that are involved in K-Ras4B - PIP2 interactions. To validate the result, we selected a subgroup of these residues for mutation if these residues are also identified as cancer associated point mutations (missense, nonsense and silent mutations) [Šolman et al., 2015]. We first examined the hydrolysis and exchange activity of K-Ras4B, including both intrinsic activity and stimulated activity by regulatory proteins p120Ras as a GTPase Activating Protein, GAP and SOS as a Guanine Exchange Factor, GEF. If the mutants are greatly altered in their hydrolysis and exchange activity, it is clear that the mutations already have effects solely on the K-Ras protein, even though they probably also affect the lipid binding of the GTPase core region.

The measured intrinsic and stimulated hydrolysis rates are shown in Fig. 7A. The K-Ras4B G12V constitutively active form was used as negative control. The activity of the other mutants were calculated relative to the wild type (the stimulated hydrolysis activity of WT with p120Ras GAP was scaled to 100). Among the 11 mutations, the K16E mutation substantially decreases (essentially disrupted) the p120 GAP stimulated GTP hydrolysis rate compared to the WT, while apart from the K104M and K165E/K167E mutations, all others the increases the p120 GAP stimulated GTP hydrolysis rate compared to the WT. The intrinsic and stimulated nucleotide exchange rates are shown in Fig. 7B. The K-Ras4B S17N mutant is a dominant negative form in vitro and was used as negative control. Among the 11 mutations, the K16E and K147E mutations substantially decrease the SOS GEF stimulated exchange rate compared to that seen with the wild type GTPase. The other mutations show little/no significant changes.

**Figure 7.**
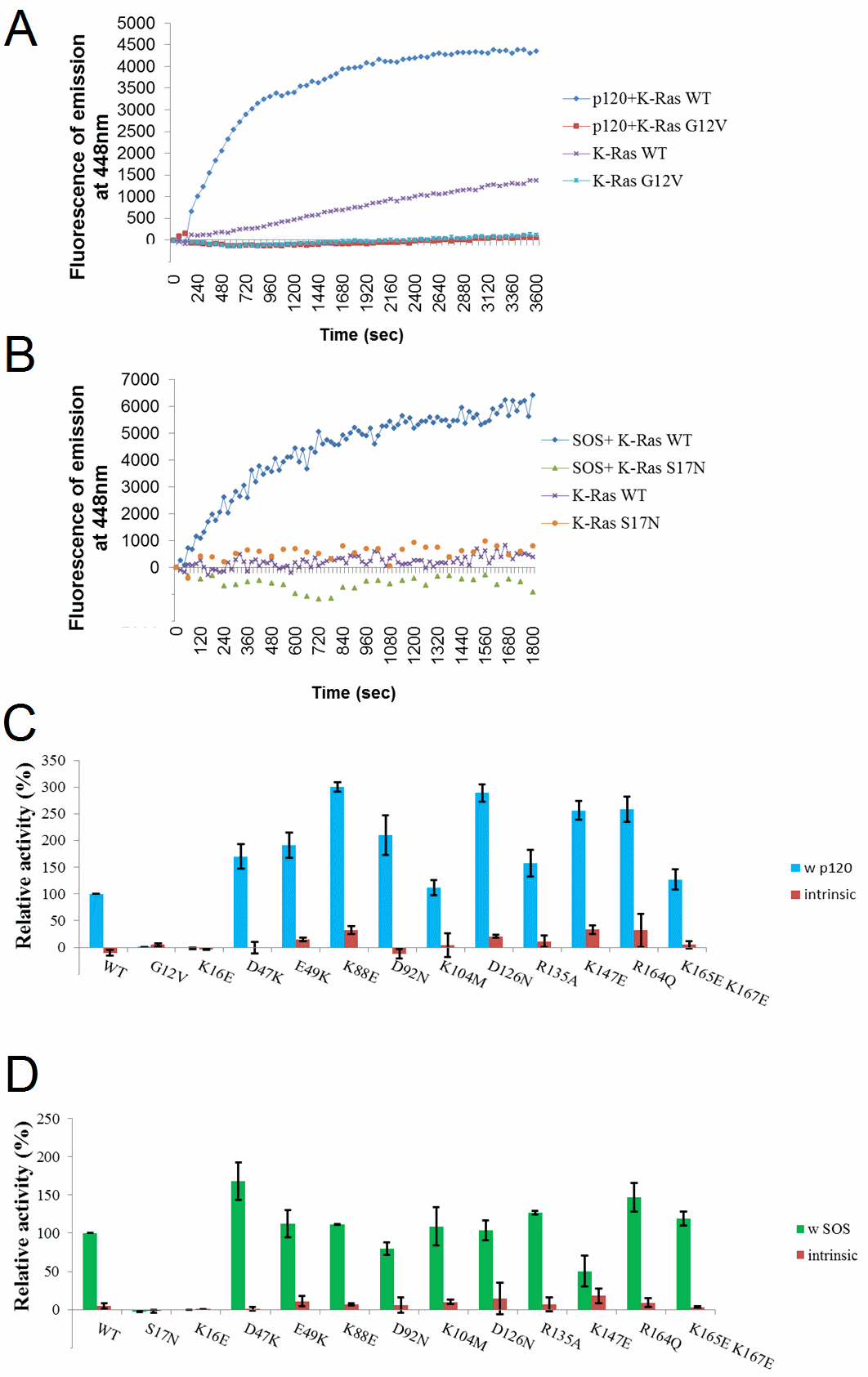
GAP and GEF activity of designed K-Ras4B mutants. Assays were carried out for the intrinsic and p120Ras GAP aided GTP hydrolysis activity and for the intrinsic and SOS GEF stimulated nucleotide exchange activity. A. Representative time-courses of intrinsic and p120 GAP stimulated GTP hydrolysis activities are shown for K-Ras WT and G12V. B. Representative time-courses of intrinsic and SOS GEF stimulated exchange activities are shown for K-Ras WT and S17N. C. Histograms of relative intrinsic and p120 GAP stimulated hydrolysis activities of K-Ras constructs, with intrinsic hydrolysis activities shown in red, stimulated hydrolysis rates shown in blue (wild type K-Ras was scaled to 100). D. Histograms of relative intrinsic and SOS GEF stimulated exchange activities of K-Ras constructs, with intrinsic exchange activities shown in red, stimulated exchange rates shown in green (wild type scaled to 100%). Note: the intrinsic rates in C. and D. and their uncertainties are plotted as multiplied by a factor of three in order to increase visibility.

### Effect of mutations on K-Ras4B transforming activity

Next we wished to study the role of lipid binding in the biological activities of the mutant K-Ras4B mutant G12V in intact cells. We employed the established ability of this mutant protein to induce the loss of contact inhibition in confluent cultured fibroblasts [Lin et al. 1999], an accepted surrogate for oncogene-induced cell transformation [Martin et al. 2001]. The cell transformation results of K-Ras4B mutants are shown in Fig. 8. Mutations of the following residues caused a significant impairment in the focus formation activity of K-Ras4B.G12V: K16, D47, D92, K104, D126, and K147. Among these, the effect of mutations K16E, D47E and K147E is not attributable to disruption of lipid binding alone, since these mutations altered the GAP or GEF activity too (see Fig. 7). By contrast, we have shown that mutation D92, D126 and K104M have essentially unaltered nucleotide hydrolysis and exchange activities. The impairment in the focus formation, thus, lends direct support for the involvement of these residues in lipid binding and that the lipid binding is essential for the protein’s biological activity.

**Figure 8:**
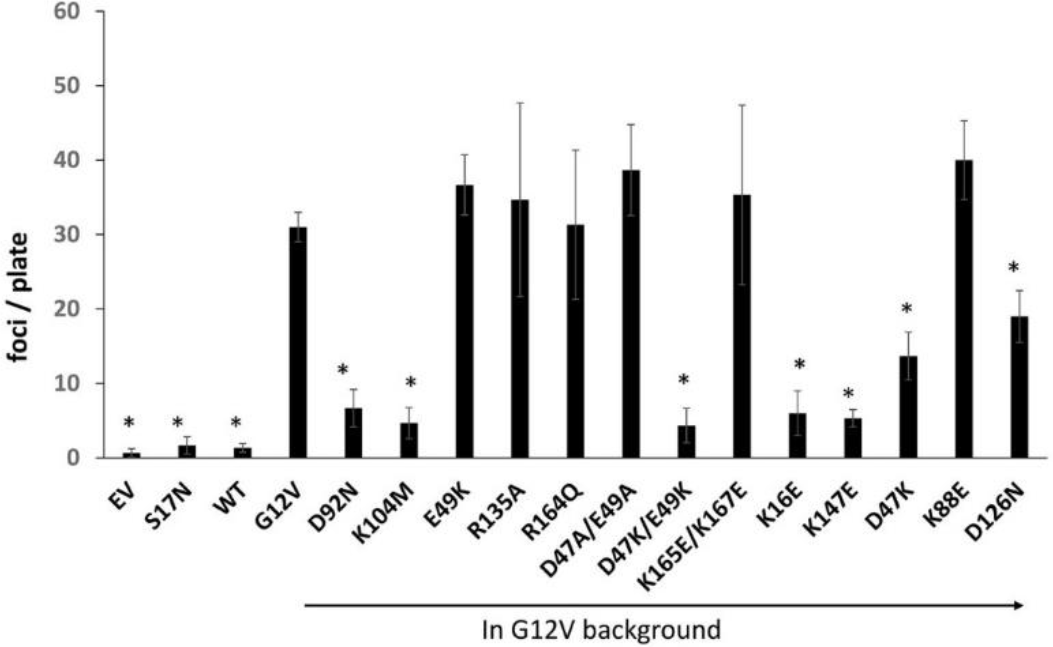
Transformation activity of designed K-Ras4B mutants as measured in foci formation of NIH3T3 cells. Y-axis of the histograms is the foci count per plate. Ev indicates untransformed cells. K-Ras4B G12V and K-Ras4B S17N were included as positive control and negative control respectively. The foci formation activity of the designed K-Ras4B mutants were all based on the oncogenic G12V background.

### Effect of mutations on intracellular localization of K-Ras4B

We further examined how mutations of lipid-binding residues affect K-Ras4B’s intracellular localization in cultured cells. Fig. 9 shows results from microscopy experiments, where transfected K-Ras4B alleles were visualized with anti-K-Ras immunofluorescence and the actin cytoskeleton highlighted with fluorescently-tagged phalloidin. As anticipated and reported earlier [van der Hoeven, 2013], we observed the localization of the constitutively active K-Ras (G12V) allele at the cells’ periphery, along with cortical actin that marks the plasma membrane. However, different mutations of lipid-binding residues have different impact on K-Ras localization. The K104M mutation completely disrupted plasma membrane localization, indicating that this residue indeed is critical in lipid binding and thus intracellular localization of K-Ras4B. This provided a direct explanation for the inactivity of this construct in the transformation assays (Fig. 8). On the contrary, the D92N and D126N alleles did not significantly affect the localization, and remained localized at cells’ periphery. But meanwhile transformation assays show that these two constructs substantially diminished the foci activity of NIH3T3 fibroblasts. The possible explanation is that these two residues indeed are involved in K-Ras-PIP2 membrane binding and tune the orientation of K-Ras at the membrane. Changes of K-Ras4B orientation on the membrane rendered K-Ras4B unable to bind certain downstream regulators, subsequently diminishing the transformation activity.

**Figure 9.**
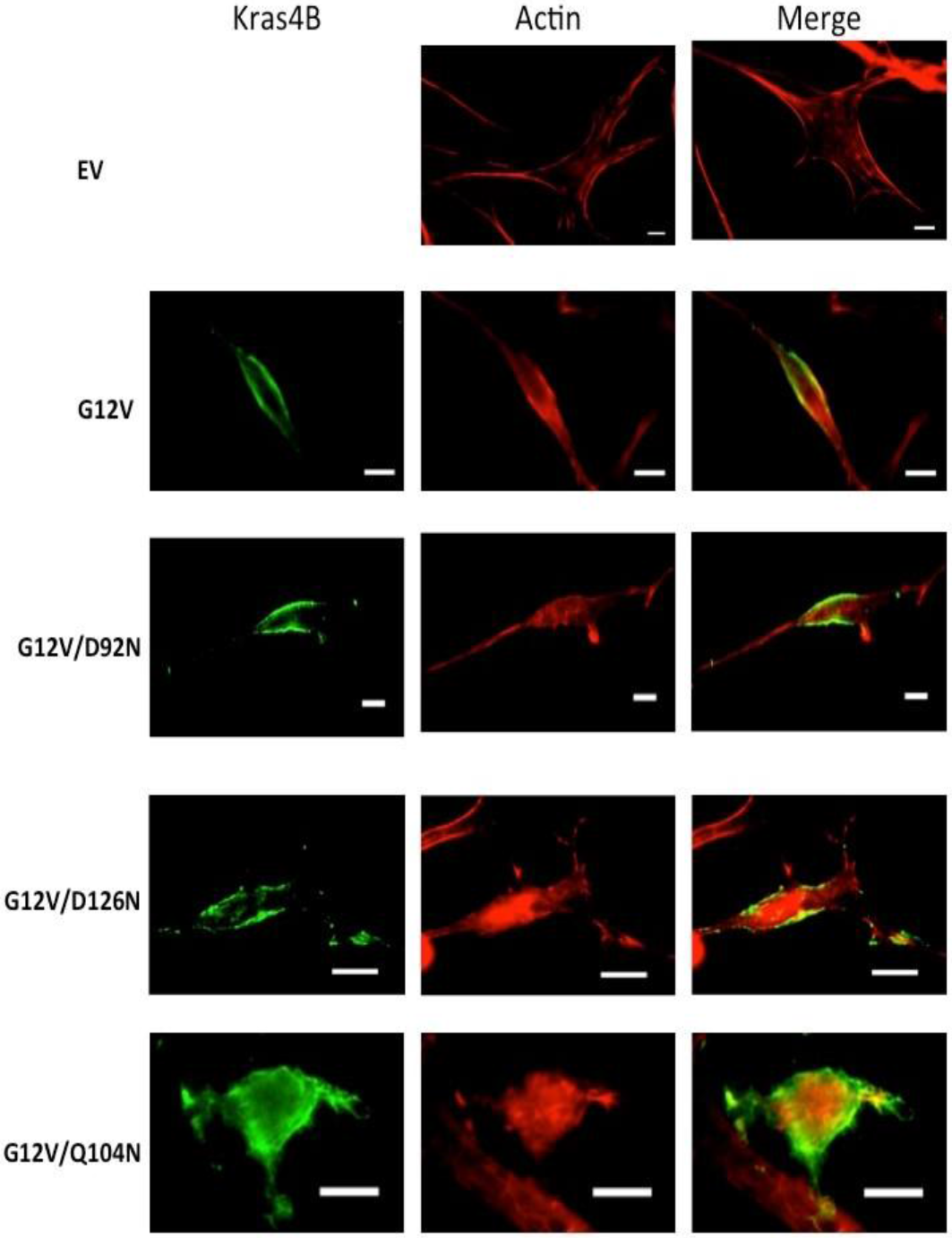
Intracellular localization of designed K-Ras4B mutants in cultured NIH3T3 cells using fluorescence microscopy. Transfected K-Ras4B alleles were visualized with anti-K-Ras immunofluorescence and the actin cytoskeleton highlighted with fluorescently-tagged phalloidin. Images are shown for Ev, K-Ras4B G12V, and designed K-Ras mutants D92N/G12V, D126N/G12V, K104M/G12V. Ev, indicates untransformed cells, and K-Ras4B G12V was included as positive control.

## Discussion

Our experimental data from a lipid strip assay, solution NMR spectroscopy, microscale thermophoresis and also all atom molecular dynamics simulations clearly show that K-Ras4B directly interacts with PIP2. Binding is more extensive and appears stronger when several protein sites are involved simultaneously, consistent with the concept of binding avidity (multivalency) [e.g. Bagnard and Smith, 2012, Banjade & Rosen, 2014]. Thus, we confirm that the interaction of K-Ras4B with PIP2 involves both the G- domain and the HVR region, as shown by experiments with K-Ras4B where the HVR was truncated. Furthermore, our NMR data and also previous simulations with K-Ras4A and PIP2, suggest that the G-domain samples multiple orientations relative to the PIP2 membrane [Li et al., 2017]. Such dynamic protein configurational states at the membrane are likely to be important for cell signaling kinetics.

Involvement of G-domain of K-Ras GTPases in binding to phosphatidylserine (PS) containing membranes have been previously reported in several papers. In a simulation study, Prakash and Gorfe et al., reported the involvement of helices 3, 4 and β-strands 1–3, as well as helix 2 on the opposite face of the catalytic domain in K-Ras4B G12D interaction with POPS/POPC bilayers [Prakash, et al., 2016]. More recently, the interaction of C-terminally membrane anchored K-Ras4B with PS was also studied experimentally by Ikura and his colleagues using nanodiscs [Mazhab-Jafari et al., 2015]. This study revealed two, if not possibly more, orientation states, but also suggested a GTPase nucleotide bound state specific shift between these states.

In general, our results for K-Ras4B at the PIP2 membrane are in good agreement with this experimental study of K-Ras4B at PS containing nanodiscs [Mazhab-Jafari et al., 2015]. While our orientation 1 (OS1) resembles the GTP state orientation, orientation 2 (OS2) resembles the GDP state orientation in their paper, there are also some differences, which likely arise because of the different nature of PIP2 compared to POPS/ PS lipid. Specifically, PIP2 has a -4 charge compared to the -1 charge of PS and also has a larger head-group. As we discussed in Li et al., 2017, the electrostatic interaction between K-Ras and PIP2 dominates the binding, and thus may account for the orientation change of K-Ras on PIP2 containing membranes (increased population of OS1-like states). However, it is worth noting that in the experimental study K-Ras was bound to the membrane via a thiol-reactive maleimide-functionalized lipid [Mazhab-Jafari et al., 2015]. By contrast, our K-Ras is not anchored to the membrane and thus our experiments report more on transient events, rather than tightly bound states or transitions between them. Thus, the similarity between the results of these two studies indicate that the transient encounters occur with orientations which are equivalent to those seen in the more persistently membrane bound G-domain. This is consistent with the observation that there no significant medium or long-range G-domain contacts with the HVR in these simulations [Prakash et al., 2016; Li et al., 2016]. Overall these findings are consistent with the emerging view that while the HVR plays a major role in lipid binding selectivity and in strengthening Ras-membrane interactions [Zhou et al., 2017], the orientation states of the G-domain are closely related to the identity (also nucleotide bound state and oncogenic mutation) of the Ras GTPase, with the HVR playing a minor role [Prakash & Gorfe, 2017].

The interactions of the G-domain with membrane imply that they may help the GTPase to associate with membranes prior to C-terminal lipidation or in cases, such as binding to the mitochondrial outer membrane, where lipidation is not needed [Sung et al., 2013]; thus our in vitro biophysical study were carried out with the full length but non-lipidated protein, or with non-lipidated HVR-truncated protein. Such interactions between the G-domain and PIP2 containing membranes could especially relevant biologically in cases where the lipid group is not yet attached, or when it is occluded by other GTPase regulators, such as phosphodiesterase, PDEδ [Suladze et al., 2014] or other lipid binding/GTPase chaperone proteins [Zhang et al., 2018]. While it is long established that lipidated HVR helps Ras GTPases anchor to the membrane, decades of research into the development of compounds that interfere with HVR lipidation, processing, and subsequent binding to membranes have failed to provide effective therapeutic reagents [Lobell et al., 2002 a, b]. Thus the binding interface between G-domain and membrane may provide a novel avenue to target K-Ras.

Electrostatic interactions play an important role in K-Ras4B binding with PIP2. As shown in the contact frequencies analysis of the simulation results, charged residues have a high frequency in contact with PIP2 membrane. But different from our intuition, not only positive but also negative residues participated in binding with the anionic PIP2 membrane. While the positively charged residues interact directly with the negatively charged PIP2 head group, the negatively charged residues were indirectly bridged with PIP2 by sodium ions. This is supported by NMR results too. Multiple residues identified by NMR in K-Ras4B binding with PIP2 are immediately adjacent to the corresponding charged residues listed in the contact frequency analysis of simulation. A similar role of negatively charged residues and bridging sodium ions was also found in our recent simulation study of K-Ras4A binding with PIP2 lipid [Li et al., 2017]. At the same time work by Hancock and colleagues have also implicated a role of hydrogen bonding of Arg vs. Lys sidechains and even for hydrophobic interactions between the HVR and lipids [Zhou et al., 2017]. Such information should be helpful in the future design of possible therapeutic agents that aim to disrupt or stabilize K-Ras interaction with membrane.

However, the interaction of K-Ras4B with a PIP2 containing membrane is dynamic rather than fixed, likely more so since-as noted- in this study K-Ras is not membrane anchored. Mentioned in the results section, we were able to exploit the sensitivity of NMR to dynamics on a range of different timescales and record PRE-data upon binding to liposomes, doped with PIP2 and spin-labeled lipid. From our results, we propose there is an equilibrium of unbound protein and multiple orientations of K-Ras4B that are bound to PIP2 membrane. This may be important for the diverse functions of K-Ras. The G-domain of Ras GTPases binds GDP/GTP and associates with effectors, GTPase exchange factors (GEFs), and GTPase activating proteins (GAPs) [Vetter et al., 2001]. The multiple orientations thus enable K-Ras interact with certain regulatory and effector proteins at different contexts, while hindering the interaction with others. Indeed, a recent nuclear magnetic resonance (NMR) study of K-Ras proposed that orientation preference of K-Ras4B at POPS bilayer membrane is nucleotide dependent [Mazhab-Jafari et al., 2015]. Specifically, the GTP state K-RAS4B binding with membrane occludes its interaction with effectors. On the contrary, GDP state K-Ras4B, as well as in the oncogenic G12D mutant, the G-domain interaction with membrane exposed its binding interface for effectors and regulatory proteins [Mazhab-Jafari et al., 2015]. Another simulation as well as experimental study showed that H-Ras sampled two major orientations on DOPC bilayer, and that the population of the two orientations can be tuned by mutating residues in the G- vs. the HVR domain [Gorfe et al., 2007]. Simulation studies on K-Ras also suggested two major orientations of K-Ras at a DOPC bilayer [Jang, 2016], a DMPC bilayer [Abankwa, et al., 2010] and at POPC/POPS bilayer [Prakash, et al., 2016]. So the multiple orientations of K-Ras4B on a PIP2 containing membrane may not be unique but are probably a common phenomenon in interactions of different GTPases with different lipids.

PIP2 binding to K-Ras4A [Li et al., 2017] and here K-Ras4B, appears to stabilize an OS1-like state (denoted O3 in our previous paper), which has the effector and regulatory binding switch regions exposed to the cytoplasm and thus is expected to increase the signaling function of the GTPase, relative to other membrane bound states which occlude these regions. How the membrane regulates the functions of K-Ras in a live cell environment, however, is rather complicated.

First, K-Ras can be postranslationally modified by phosphorylation, nitrosylated, ubiquitinated or acetylated on certain residues, all of which may affect GTPase function and targeting. In the case of the mutation of K104, this will affect a primary acetylation site. Yang et al., (2012) showed that the K104Q mutation (an acetylation mimic) suppressed transformation activity (similarly to K104M in our case) but by contrast to our results did not affect localization to the plasma membrane. Another more recent study found K104Q compromised both GEF and GAP activity, but had no effect on cell transformation, possibly due to compensatory mechanisms in the cell [Yin et al., 2017]. Our results are different in that K104M has normal GEF and GAP function, but is compromised in its localization and transformation ability. The former results may be explained by methionine sidechain being of similar geometry to lysine, in our case, while the latter, may indeed also be due to a difference in residue sidechain character, with glutamine still being able to hydrogen-bond and interact with PIP2, whereas a methionine may not be.

Second, membranes are a complex mixture of several lipids [e.g. Yang et al., 2018] and local clustering of lipids (and the involvement of proteins in this process) is just beginning to be understood [Banjade and Rosen, 2014; Lu & Fairn, 2018]. For example, it may be expected that the PS can, to some extent, mask, if not compete with protein - PIP2 interactions in physiological membranes. However, this could be protein specific as indicated by one recent study, where the apparent K-Ras4B lipid interaction specificity could be switched by a lysine to arginine mutation or Ser181 phosphorylation in the HVR region [Zhou et al., 2017].

Third, several studies have provided evidence for the existence and biological relevance of Ras dimers, establishing a new mechanism for regulating Ras activity [e.g. Inouye et al., 2000, Güldenhaupt et al., 2012]. Research by the Hancock group and others have shown that N-, K- and H-Ras assemble into higher order oligomers and nanoclusters, and do so in an isoform-specific manner [Muratcioglu et al., 2015; Nan et al., 2015; Plowman et al., 2005; Zhou and Hancock, 2015]. However, recently, neither K-Ras dimerization, nor clustering could be shown on supported lipid bilayers with a range of lipid compositions, suggesting that additional proteins, e.g. the cytoskeleton as a scaffold, may be required [Chung et al., 2018]. The situation in cancer cell can be even more complicated, due to the different types of K-Ras mutations [recently Ambrogio et al., 2018], and mutations/abnormal expressions of Ras regulating proteins and signaling lipids too.

To resolve the mechanism of Ras regulation by membranes, further experimental and computational studies of Ras cancer mutants, of post-translationally modified forms, of Ras, of Ras in complex with regulatory proteins, of dimers and oligomers of K-Ras, -all at the membrane-will be needed. In addition, most effector proteins bind to the Ras GTPases at the membrane. For example, our lab has recently published a simulation study of K-Ras - C Raf (CRD-RBD domain) complexes at a POPS membrane [Li et al., 2018], which illustrated the complexity of such systems when the effector protein domains interact with the membrane as well.

## Concluding Summary

The present data provide, to our knowledge, the first experimental study of Ras-membrane interaction using liposomes and NMR, and also provide a residue level structural report of the K-Ras G-domain binding to PIP2. Binding to PIP2 likely provides an additional mechanism to tune, if not regulate GTPase signaling activity in cells.

## Experimental procedures and methods

### Protein constructs, expression, purification and site directed mutagenesis

The cDNA for human K-Ras4B (residues 1-188) was obtained from cDNA.org (hosted at Bloomsburg University) and subcloned into pET28a using Nde1 and BamH1 restriction sites. HVR truncated K-Ras4B (residues 1-169) and the mutants were made using the QuickChange Lightning site directed mutagenesis kit (Agilent). A subgroup of residues that are identified in K-Ras - PIP2 membrane binding were selected for mutation, if these residues are also identified as cancer associated point mutations (missense, nonsense and silent mutations) [Šolman et al., 2015]. Transformed *E. Coli* strain BL21 (DE3) bacteria (Novagen) were grown at 37°C in LB, and induced with 1 mM IPTG for expression at 25°C overnight. Bacterial pellets were resuspended in the following buffer: 20 mM Tris-HCl, 150 mM NaCl, 0.5 mM TCEP (tris(2-carboxyethyl)phosphine hydrochloride), 4.0 mM MgCl_2_, pH 7.5, with added protease inhibitors (PMSF 1 mM, bezamidine 10 mM, leupeptin 42 μM, antipain 3 μM). Following sonication, lysates were clarified by centrifugation and recombinant proteins were purified from the supernatant using Ni-NTA agarose (Qiagen). Purified K-Ras proteins were dialyzed against NMR buffer (20 mM Tris-HCl, 20 mM NaCl, 1 mM TCEP, and 4 mM MgCl2, pH 7.4). Purified proteins were >90% pure identified by SDS-PAGE. For NMR experiments, K-Ras was ^15^N uniformly labeled by growing bacteria in M9 medium containing ^15^NH_4_Cl as the sole nitrogen source.

Similar to other protocols, K-Ras was loaded with nucleotides (GTP or its non-hydrolyzable analog GMPPNP) by incubating 100 μM freshly purified protein with 0.5 mM nucleotide in 20 mM Tris-HCl, pH 7.5, 100 mM NaCl, 5 mM EDTA, 2 mM DTT at 30°C for 10 min. The reaction was stopped by adding 50mM MgCl2. A PD-10 Sephadex G-25 desalting column was used to remove the excess nucleotide, and to change the buffer to 20mM Tris-HCl, pH 7.5, 150mM NaCl, 1mM TCEP, 4mM MgCl_2_. The catalytic domain of p120 RasGAP (a gift from Dr. R. Ahmadian) subcloned into pET28 and expressed and purified as described previously [Ahmadian et al., 1996]. The SOS catalytic domain in a pProExHTb vector was gift from Dr. J. Kuriyan. This protein was expressed and purified as described [Sondermann et al., 2004]. Proteins were placed at 4 °C/on ice for immediate use or flash frozen with additional 5% glycerol and stored at −80 °C.

### Protein-lipid overlay assay

Nitrocellulose lipid strips (Echelon Biosciences) were blocked with 3% fatty acid-free BSA in PBS / 0.1% Tween20 for 1 hr, and incubated for 1 more hr with 100 nM mouse purified His-tagged K-Ras at 4 °C. The membranes were then washed four times in PBS buffer. After washing, membranes were incubated with monoclonal anti-His6 antibody (ThermoScientific) at 1:2000 dilution for 1 hr at 4 °C, followed by additional washing and incubation with Goat anti-mouse-IgG-horseradish peroxidase-conjugated antibody (Santa Cruz) at 1:2000. After final washing, membrane-bound K-Ras was visualized by chemi-luminescence (Sigma) [Dowler et al., 2002].

### Liposome preparation

All lipids and lipid head groups were purchased (Avanti Polar Lipids Inc). For the full length lipids, as a first step, the lipid-chloroform solutions were dried under nitrogen gas, and then hydrated in 20 mM Tris-HCl, pH 7.4, 20 mM NaCl, 2 mM TCEP, and 4 mM MgCl_2_. For MST experiments, fluorescent labeled lipids 1,2-dioleoyl-sn-glycero-3-phosphocholine (DOPC), TopFluor^®^ PIP2 (3, 5), TopFluor^®^ PIP2 (4,5), and TopFluor PIP3 (3,4,5) were mixed at the stated ratios. Rehydrated lipids were extruded through polycarbonate membranes with 100 nm pore size (Avestin) using mini extruder and following the manufacturers’ manual (Avanti Polar Lipids Inc) to a final lipid concentration of 500 μM. The diameter of the prepared liposome was determined by dynamic light scattering to be 120 nm +/− 20nm.

### Microscale thermophoresis (MST)

MST was measured with fluorescently labeled liposomes to a final concentration of 5 nM, titrated by a serial dilution of K-Ras (294 μM to 0.1436 μM). K-Ras was tested in NMR buffer (see below), supplemented with 2 mg/ml BSA. Alternatively, the K-Ras protein was fluorescently labeled with dye NT-647, using the Monolith Protein Labeling Kit RED-NHS (NanoTemper Technologies). Mixtures of the protein-lyposome solutions were filled into hydrophobic glass capillaries (Nanotemper Technologies) and were measured with a Nanotemper Monolith NT.115 system (75% light-emitting diode, 20% IR laser power). The protein-lyposome dissociation constant (*K*_*d*_) was obtained by fitting the binding curve with the quadratic solution for the formation of a protein-liposome complex (assuming 1:1 binding), calculated from the equation: [PT] = 1/2*(([P0] + [T0] +*K*_*d*_ -(([P0] + [T0] + *K*_*d*_)^2^ - 4*[P0]*[T0])^1/2^)

Where [P0] is the concentration of the total fluorescent K-Ras, [T0] is the total lipid concentration. [PT] is the concentration of the formed protein-lipid complex. The concentration of the complex ([PT]) is derived from the fraction of fluorescent molecules that formed the complex x = [PT]/[P0], in which x directly corresponds to the signal obtained in the MST measurement (normalized fluorescence).

### Preparation of Nanodiscs and Liposomes for NMR

The MSP1D1 protein was used to make nanodiscs [e.g. following Puthenveetil et. al., 2017]. The pGBHPS-MSP vector encoding membrane scaffold protein (MSP) variant 1D1 was obtained from AddGene. MSP1D1 was expressed and purified according to established protocols [Kucharska et. al., 2015; Denisov et. al., 2004]. Briefly, MSP1D1 was expressed in *E. Coli* (BL21) grown in Terrific Broth, and 37 °C and when OD600 reached 2.0 protein expression was induced with 1 mM IPTG for 1 hr followed by a 2.5 hrs incubation at 28°C. MSP1D1 was purified using Ni-NTA agarose (Qiagen) resin, and His tag was cleaved with TEV protease and further purified as described elsewhere [Ritchie et al., 2009; Mazhab-Jafari et al., 2013]. The efficiency of cleavage was monitored using SDS-PAGE to be > 90%. For the nanodisc preparation, PIs, PE-DTPA and DOPC were mixed with at a molar ratio of 10:5:85. The lipid solution was dried under nitrogen, followed by drying under high vacuum for at least 4 h to remove the residual organic solvent. The dry film was solubilized in 100 mM cholate in the NMR buffer (20 mM Tris-HCl, 20 mM NaCl, 2 mM TCEP, and 4 mM MgCl2 and 0.01% NaN3, pH 7.4) to a final phospholipid concentration of 36 mM. The solution was subjected to three freeze/thaw cycles, vortexed, and sonicated to clarity in an ultrasonic bath. MSPD1 was added to this lipid solution at a lipid:protein molar ratio at 80:1. The lipid-protein mixture was incubated for 1h with gentle rotation at 20°C. Immediately following the incubation, sodium cholate was removed from the MSPD1-lipid mixture by three sequential dialyses against NMR buffer. After the dialysis step, nanodiscs were further purified by size exclusion chromatography in NMR buffer on Superdex 200 10/300 (GE Healthcare). Finally, nanodiscs were concentrated using Amicon centrifugal units of 10 kDa MWCO to the desired concentration of approximately 250 μM for NMR experiments. The nanodiscs were measured with dynamic light scattering (DLS) and found to have an average diameter of 10 nm. +/-1.8nm.

For the NMR experiments that detect binding via Paramagnetic Relaxation Enhancement (PRE), nanodiscs and liposomes were prepared as above but were supplemented with 3.5% 1,2-distearoyl-snglycero-3-phosphoethanolamine-N-diethylenetriaminepentaacetic acid [gadolinium salt; PE-DTPA (Gd^3+^)]. The DOPC, PI(4,5)P2, PI(3,5)P2, PI(3,4,5)P3 as well as PE-DTPA (Gd^3+^) lipids were purchased from Avanti Polar Lipids. In NMR experiments, four kinds of samples were used: DOPC; DOPC+ PE-DTPA (Gd^3+^) (96.5: 3.5); DOPC+ PIP2 (90: 10); and DOPC+ PIP2+ PE-DTPA (Gd^3+^) (86.5: 10:3.5). Data from NMR experiments carried out with DOPC liposome and DOPC+ PE-DTPA (Gd^3+^) were used as a control, validating that there is no significant interaction of K-Ras with nanodiscs or liposomes which consisted of DOPC lipid only (i.e. have no PIP2 component).

### NMR spectroscopy

NMR spectra for K-Ras interaction with lipids using lipid head groups and nanodiscs were recorded at 25 °C on a Bruker Avance II 800 MHz spectrometer equipped with a TXI cryoprobe. To improve the spectra quality, the NMR spectra for the K-Ras interaction with lipids using liposomes were recorded at 37°C, yielding sharper lines and apparently, slightly more dispersed spectra. All NMR measurements were carried out in NMR buffer (see above). For experiments using the lipid head group IP3(1,4,5), the concentration of K-Ras protein was 150 μM, and the protein:IP3 ratio was to 1:3. In NMR experiments using nanodiscs as membrane mimetic, the concentration of K-Ras4B protein was 80 μM, the ratio of protein to nanodiscs was 1:1.2 (ratio of protein to total lipids is 1:96). In NMR experiments using liposomes, the concentration of K-Ras4B protein was 80 μM, the ratio of protein to the total lipid concentration of the liposomes was 1:100 (with 10 lipids as PIP2). The spectra were collected with 1024*180 (F1*F2) points and 32, 80, 96 scans and were processed with NMRPipe [Delaglio et al., 1995] and analyzed with software Sparky (Goddard TD & Kneller DG, SPARKY 3, University of California, San Francisco).

### Analysis of line broadening and chemical shift perturbation (CSP) for NMR experiments

For PRE measurements, the peak intensities in the spectrum of K-Ras in presence of nanodiscs or liposomes containing PE-DTPA (Gd^3+^), were compared with those of a control sample prepared without spin label. The cross-peak intensities were measured using Sparky by Gaussian line fitting. Any small difference in protein concentration between samples was corrected by normalization of the calculated intensity ratios against the highest observed I*/Io (where I* is the peak intensities in the spectrum of K-Ras in presence of nanodiscs or liposomes incorporating 3.5% PE-DTPA (Gd^3+^), and Io is that in the paramagnetic ion-free nanodiscs or liposomes). The weighted average chemical shift perturbation (CSP) were calculated as: Δδavg = [(ΔδH)^2^ + (ΔδN/5)^2^/2]^0.5^. NMR assignment of small GTPases are typically challenging projects [e.g. Cao et al., 2013]. Thus, assignments were transferred, with a reasonable level of confidence for dispersed peaks, from the published NMR assignment for human K-Ras (residues 1-166) in the GDP-bound form at a physiological pH of 7.4 [Vo et al., 2013]. Peaks which could not be assigned this way were followed as unassigned (UNA) and are nevertheless informative. All peak intensities were given uncertainties relative to spectral baseline noise and those uncertainties were propagated to the peak intensity change ratios plotted.

### Molecular dynamics simulations

A membrane consisting of 284 POPC and 16 PIP2 lipid molecules was generated by the program/website CHARMM-GUI [Jo et al., 2008]. The membrane was equilibrated for 100 ns at 310K in solvent with counterions. Simulations were carried out with full length K-Ras, which was adopted from a previous publication (Prakash et al., 2016b, Li et al., 2017) and was originally built by ligating the crystal structure of G12D K-Ras (PDB: 4DSO, residues 1-173) to a K-Ras4B HVR and lipid anchor (tK, residues 174-185). This structure was chosen because it extends the C-terminal helix to res. 173 and for the results to be comparable to other simulations. The parameter for the farnesyl group at the C-terminal of HVR region was produced by the CHARMM generalized force field (CGenFF) [Vanommeslaeghe et al., 2012]. Previously, Prakash et al. showed that K-Ras4B G12D populates two major orientations relative to the membrane (Orientation State 1, OS1 and Orientation State 2, OS2) [Prakash et al., 2016 a, b]. These were provided to us and adopted here as the initial configurations. As before, the farnesyl group was pre-inserted into the membrane [Li et al., 2017]. The CHARMM36 force field including the CMAP correction was applied to the system [Huang, et al., 2013, Buck et al., 2006]. The TIP3P model was used for water.

The system was neutralized and provided a near-physiological ion concentration of 0.15 M NaCl. In the all-atom simulations, the electrostatic interaction was treated by the Particle-Mesh Ewald (PME) method. The van der Waals interaction was cut at 1.2 nm. The time step was set as 2 fs. Temperature was coupled by using Langevin thermostat at 310 K, whereas pressure was 1 bar controlled by the semi-isotropic Langevin scheme. All these systems ran for the first 30 ns using the NAMD/2.10 package [Phillips et al., 2005], and then were transferred to the Anton supercomputer [Shaw et al., 2009] for another 350 ns of simulation.

### GTP hydrolysis assays

The kinetics of GTP hydrolysis were measured by monitoring the release of inorganic phosphate using a fluorescently labeled, phosphate-binding protein (PBP) sensor, as previously described [Sosa et al., 2009]. 2μM of K-Ras.GTP (loaded as described above) was incubated with 2 μM MDCC-phospho-binding proteins (PBP) (Thermo Scientific) until the baseline was stabilized (2-3 min). 75 nM p120-GAP was added to the reaction. Released Pi was monitored as changes in the fluorescence over time. The fluorescence was measured with Excitation at 425 nm (5 nm band width) and emission at 465 nm (7 nm band width) in a Tecan Infinite M1000 microplate reader. All experiments were done in triplicates, and data shown represent averages and standard deviations. Reported values derive from the initial velocity of the hydrolysis reaction (fluorescence change over the first 5 minutes after mixing), and the relative activity (fold-difference between initial velocity with vs. without p120-GAP, or mutants vs. wild-type K-Ras).

### Nucleotide exchange assays

Nucleotide exchange reaction was measured by monitoring the time-dependent fluorescence change of Mant-GDP as it occupies the Ras nucleotide binding pocket [Leonard et al., 1994]. 2μM of K-Ras.GTP was incubated with 1.25μM mant-GDP (Jena Bioscience) in 20mM HEPES, 50mM NaCl, 4mM MgCl_2_, 0.5mM TCEP, pH 7.5 at 25°C for 5 min for the base line to stabilize. 200nM SOS was added to the reaction and GEF activity was monitored by the fluorescence change of mant-GDP (excitation: 355nm, emission 448nm) over 30 mins in a Tecan Infinite M1000 plate reader at 25ΰC. Again the initial rates were used for assessment of the exchange kinetics. All experiments were done in triplicates. Shown data are averages and standard deviations. For each protein the initial rate and rate in the control were calculated (initial rate with vs without SOS, or mutants vs. wildtype K-Ras).

### Focus formation in cell assay

Approx. 250,000 NIH3T3 cells were seeded in triplicate in 35 mm dishes in DMEM containing 10% calf serum. The next day, the cells were transfected with 250 ng of the indicated K-Ras allele cDNA in the pCEFVL plasmid or together with 3 μg of empty pCEFVL vector using Polyethylenimine (PEI; Polysciences Inc.) according to published protocol [Durocher et al., 2002] Transfection mixture contained 3 μL PEI: 1 μg DNA per well in serum-free media. After 24 hours, each well was transferred into triplicate 10 cm dishes and grown in DMEM containing 2% calf serum for two weeks. The media was replaced every 2 days. After washing with PBS, cells were fixed with 3.7% paraformaldehyde for 15 minutes, stained with crystal violet, and visible foci (>2 mm) were manually counted under a microscope. Values reported are averages of triplicate transfections and associated standard deviations.

### Fluorescence microscopy

NIH3T3 cells were grown on collagen-coated cover slips, transfected with the indicated K-Ras constructed using PEI as described above, and cultured in serum-free media for 17 hours. After washing, cells were fixed in 3.7% paraformaldehyde, they were permeabilized with 0.2% Triton-X100, Blocked with 2% BSA and stained with anti-K-Ras antibody (clone 3B10-2F2, Abnova). Actin cytoskeleton was stained with AlexaFluor 555-conjugated Phalloidin. Slides were imaged on a Keyence BZ-X 700 fluorescence microscope.

## Acknowledgements

**Acknowledgements:** We are grateful to Dr. J. Kuriyan (UC Berkley) for the generous gift of a bacterial expression vector of the human Sos GEF and Dr. R. Ahmadian (Univ. of Düsseldorf). This study was supported by NIGMS grant R01GM112491 to the Buck lab, and Dr. S.Cao was supported in part by grant R01GM099775 (D. Altschuler, PI). We thank to the Ohio Super Computer center for computational resources. Anton2 Computer time was provided by the Pittsburgh Supercomputing Center (PSC) through Grant R01GM116961 from the National Institutes of Health. A refrigerated centrifuge used for this project was in part funded by a donation in memory of Daniel Sheehan by the Sheehan Family Trust.

**Conflict of interest:** The authors declare that they have no conflicts of interest with the contents of this article.

## FOOTNOTES

**Author contributions:** SC1, SC2, ZLL, SJK, DM and MB conceived and designed the study; SC1 performed the NMR and MST experiments; SC2 and DM the cell biology experiments SJK carried out molecular biology, protein purification as well as GEF and GAP assays; ZLL performed and analyzed the simulations; all authors analyzed the data and wrote the manuscript.

